# A simple model of population dynamics with beneficial and harmful interaction networks for empirical applications

**DOI:** 10.1101/2024.10.15.618620

**Authors:** Malyon D. Bimler, Luz Valerie Pascal, Matthew P. Adams, Chris Baker

## Abstract

1. Population dynamic models can forecast changes in the abundances of multiple interconnected species, which makes them potentially powerful tools for managing ecological communities, yet they remain largely under-utilised in applied settings. High data requirements and the ability to only model a narrow range of ecological interactions and/or trophic levels together limits their usefulness when faced with complex and data-poor systems, where beneficial (e.g. mutualism) and harmful (e.g. competition) interactions may operate simultaneously within and between species.
2. We present a model of population dynamics that can describe a wide range of ecological interaction outcomes with a simple, unified structure. Species growth rates are constrained by a maximum growth rate parameter which prevents the risk of population explosions even in the case of mutualism. Species interactions are defined by two, not mutually-exclusive interactions matrices that describe the effects of beneficial and harmful interactions respectively, together providing the potential for the net effect of interactions between one species and another to switch from beneficial to harmful as population density increases.
3. This model recreates classic dynamics in two-species mutualistic, competitive, and predator-prey scenarios, allowing us to model a wide range of trophic levels and interaction types together within the same equation. The maximum growth rate parameter, theoretically based in intrinsic constraints on reproduction, can be parameterised from a wide range of sources including natural history, historical data, and breeding programs. We illustrate the potential of this model with a data-poor case study of a threatened species and two interacting predators.
4. This new model is generaliseable to a wide range of natural ecological communities. Its model structure lowers data requirements whilst remaining intuitive and biologically realistic, making it an accessible option for predicting community-wide population changes in applied contexts where data is sparse and/or uncertain.

## 1 Introduction

Applied ecologists and environmentalists are faced with many challenges when managing species and ecosystems at risk. Population dynamic models, which can track changes in species abundances through time, can help guide management decisions (Adams et al. 2020). These types of models describe the reciprocal effects of species on one another when they are connected through ecological interactions such as predation, parasitism, competition or mutualism. Population dynamic models are potentially powerful tools for forecasting how species respond to changes in the density of other species, allowing ecologists to predict how specific populations and ecological communities might respond to management interventions such as culling (Rendall et al. 2021), species introductions (Baker et al. 2016), and species eradication (Bode et al. 2015). Unfortunately, applying these models effectively remains challenging (Lubiana Botelho et al. 2024). Ecosystems are complex and contain species spanning multiple levels of the trophic food web and connected by a multitude of interaction types beyond that of predator and prey (Kéfi et al. 2012; Kéfi et al. 2015). These non-trophic interactions can have effects (e.g. beneficial or harmful) that are often variable and context-dependent (Saavedra et al. 2016; Ushio et al. 2018; Bimler et al. 2024), making a ‘one-equation-fits-all’ approach difficult. Moreover, data with which to parameterise these models is often sparse and biased (Reddy and Dávalos 2003), yet setting parameter values which are biologically realistic remains crucial if the output of these models are to be trusted. Population dynamic models useful to empiricists must thus prioritise a number of characteristics: they must make biologically reasonable predictions, be easy to calibrate with potentially little information, and remain intuitive enough that they are relatively easy to apply.

The vast majority of population dynamic models in use today trace their roots back to the historical Lotka-Volterra predator-prey equations (Lotka 1926; Volterra 1926), which have since been adapted into more generalised forms known as generalised Lotka-Volterra models (GLVMs, Murray 2002). GLVMs are defined by two key features: a vector of species-specific intrinsic growth rates, which describes all species’ population growth in the absence of other species, and an interaction matrix, which describes the linear functional responses of all species’ populations to the density of all other species they interact with.

GLVMs have mathematical properties which makes them incredibly useful to theoreticians (primarily linearisation at equilibrium, Schoener 1976) but suffer from various drawbacks, especially around their treatment of species interactions, as follows. In GLVMs, the effect of each interaction on a species is described by a single and constant density-dependent linear parameter. This means that interactions between species pairs (which have two parameters representing the effects on both species) are fixed and cannot vary in strength or type between contexts, yielding a rather simplified representation of community interactions. This model structure also poses a serious challenge for modelling beneficial interactions such as facilitation or mutualism, which can induce runaway population growth when there are no other factors limiting the densities of the species involved. Variations on GLVMs have been developed to account for beneficial interactions without risking population blow-ups (Dean 1983; Wright 1989; Holland and Deangelis 2010; García-Algarra et al. 2014), typically by adding parameters that limit the benefits of mutualism at high densities, but those models are predominantly reserved for mutualist systems (e.g. plant-pollinator communities, Moore et al. 2018). Likewise, GLVM variants have also been developed to account for other types of ecological interactions such as intraguild predation or higher-order interactions (Holt and Polis 1997; Mayfield and Stouffer 2017) as well as specific ecosystems (e.g. Beverton and Holt 1957). Consequently, modeling variable interactions, multiple interaction types or multiple trophic levels together can require stitching together different equations for each species or interaction type (e.g. Thébault and Fontaine 2010; Gakkhar and Gupta 2016; Mitani and Mougi 2017), requiring further investment into model construction and analysis.

Ecologists have long recognised the need to develop alternative models of population dynamics which do not suffer the same restrictions as GLVMs (Schoener 1976), and that remain intuitive when applied to a wider range of ecosystems, trophic levels and interactions types. Here we propose a model of population dynamics where model construction was guided by three priorities: (1) ability to describe a wide range of interaction outcomes, (2) parameterisation requiring little data, and (3) intuitive enough for non-specialists to apply. In our model, populations are defined by a maximum growth rate that limits how fast populations can grow regardless of their interactions, and whose value can be informed by natural history knowledge, historical data and/or literature searches in the absence of field data. Our model introduces two interaction matrices to describe beneficial and harmful interactions respectively, without the risk of population explosions. If desired, this choice of construction also allows species to switch from having beneficial to harmful effects as density increases, making our model flexible to some degree of context-dependency in species interactions. Together, these features permit connection between multiple interaction types and trophic levels within the same model equation, maintaining biological realism without sacrificing model simplicity. We first describe the model and give the equations for the single species and multispecies equilibria. We show that this model can simulate the classic population dynamics of mutualism, competition and predator-prey systems. Finally, we apply the model to a data-poor case study of an invaded ecosystem to illustrate the biological meaning of the parameters and how to estimate these parameters when information is limited.

## 2 Model

We present a new generaliseable model of population dynamics where changes in the abundance of focal species are described by: (1) a maximum intrinsic growth rate *R* which limits how fast a population can grow, (2) beneficial interactions *α* which bring a population closer to its maximum growth rate but can never exceed it, and (3) harmful interactions *β* which lower the overall growth rate. Interactions may occur with any number of species, both conspecific and heterospecific. We set *S* as the total number of interacting species *i, j*, with *i, j* ∈ {1, .., *S*}. The model is as follows:

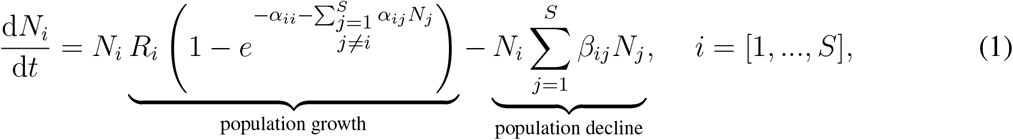

where *N*_*i*_ is abundance and *R*_*i*_ is the maximum intrinsic growth rate for each species *i*. Interaction parameters *α*_*ii*_ and *α*_*ij*_ are per-capita beneficial interactions with conspecific and heterospecific populations respectively, whilst *β*_*ii*_ and *β*_*ij*_ are per-capita harmful interactions. All parameters *R*_*i*_, *α*_*i\**_, and *β*_*i\**_ must contain non-negative entries only, where * indicates any species belonging to *S*.

Our model does not explicitly describe population growth as a function of birth, death, and dispersal processes. Instead, changes in growth rate are driven by phenomenological interactions which modulate how fast a population is growing in the first term of Equation 1 and how fast a population is declining in the second term of Eq. 1. Effectively, interactions with other organisms (conspecific or not) regulate the addition of individuals by affecting processes such as reproduction, consumption, growth, survival and colonisation. Likewise, interactions also act on the removal of individuals from a population, for example through death, competition, and dispersal. Interactions which increase the focal population growth rate (*α*_*ii*_ and *α*_*ij*_) are described with a different mathematical relationship to those interactions which decrease the focal population growth rate (*β*_*ii*_ and *β*_*ij*_). As a result, these two processes act simultaneously and do not negate one another, such that a specific interaction partner can have both beneficial and harmful effects on the focal population, while the overall net effect depends on its density (see Section 2.2).

The maximum growth rate *R*_*i*_ sets an upper limit on density-dependent growth. It encapsulates population-specific biological and environmental constraints on reproduction which are not mediated by interactions with individuals of any species in *S*. All organisms have, at the very least, metabolic and energetic constraints on how many new individuals they can produce; for example, maximum clutch size in birds, maximum seed set in flowering plants, maximum cell growth rate in bacteria. Additionally, populations in non-laboratory settings often have access to limited abiotic resources, or experience non-ideal environmental conditions which may further limit their capacity to reproduce. As a result of these intrinsic restrictions on reproduction, a population’s growth rate remains limited even in the most optimal of conditions. As biotic conditions improve (when 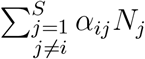 increases and 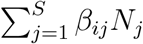 decreases), population growth will tend closer to the theoretical maximum set by *R*_*i*_.

Note that a positive maximum growth rate (*R*_*i*_ *>* 0) is not sufficient to increase population growth as it solely sets a maximum limit: it is the *α*_*ii*_ and *α*_*ij*_ parameters which determine how much of the maximum growth rate is realised. Strong beneficial interactions will bring the overall population growth rate closer to the maximum value set by *R*_*i*_*N*_*i*_, whereas weak beneficial interactions will lower the realised rate of change far more. Consequently either the *α*_*ii*_ or the *α*_*ij*_ parameters must include non-zero entries to ensure 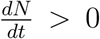, as achieving a positive population growth requires the presence of either conspecific or heterospecific individuals, respectively. We acknowledge this model structure presents a philosophical departure from how many traditional models of population dynamics (including GLVMs) conceptualise intrinsic growth rate and ecological interactions, and hope the benefits of our choice become evident upon further reading. In terms of biological justification for our model structure, few (if any) organisms are capable of thriving as completely isolated individual units: many species require the presence of other individuals for sexual reproduction, or benefit from aggregation (Bertness 1989) and other forms of cooperation (Bronstein and Sridhar 2023). Moreover, and without depending on other organisms as a food source, primary producers still benefit from the presence of other organisms to prepare or improve environmental conditions (Bruno and Bertness 2000), produce metabolites that are inaccessible or difficult to synthesize (Morris et al. 2012), and maintain the proper functioning of Earth’s biogeo-chemical cycles (Lovelock and Margulis 1974).

Including the *α*_*i*._ terms within an inverse exponent sets a saturating limit to their positive effect on the growth of population *i*, as the sum of the exponent is bounded between 1 (when 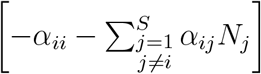 → 0) and 0 (when 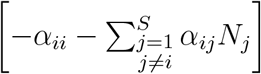 → −∞). Note that *α*_*ii*_ and *α*_*ij*_ have different units, as *α*_*ij*_ is multiplied by population density *N*_*j*_ whereas *α*_*ii*_ is not. The mech-anisms behind *α*_*ii*_ can include, but are not limited to, increases in reproductive output when there are more mates to choose from, increased defense or evasion from consumers, and ameliorations to the physical environment. Parameters *α*_*ij*_ on the other hand accounts for increases in growth rate or survival due to facilitative interactions with other populations or species, whether mutulistic, commensal, or asymmetric (e.g. benefits to consumers from predation).

Lastly, harmful interactions with both conspecific and heterospecific populations are captured by the 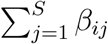 terms which lower the population growth rate of the focal population *i*. These 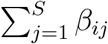 parameters describe ecological interactions which lower the overall growth rate of the focal species, such as competition, predation (including omnivory), and parasitism.

### 2.1 Equilibrium solution for a single population

In the case of a single species *i* which does not interact with any other species, population growth follows a sigmoidal model (e.g. Simpson et al. 2022; Mills et al. 2024) of the form:

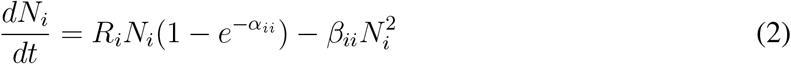

We can identify the equilibrium abundance 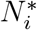 in the case of single population by setting the left-hand side of Eq. 2 equal to 0, and arrive at:

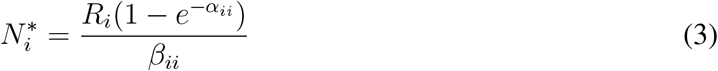

which is also the carrying capacity of this one-species system. Equation 3 illustrates a key feature of the model: in the case of a single population, carrying capacity is highly dependant on the strength of intraspecific facilitation *α*_*ii*_ and intraspecific competition *β*_*ii*_. These two terms must be positive for a species population to be able to grow in isolation and reach a positive equilibrium in isolation, respectively. That *α*_*ii*_ must be positive means our model assumes that for a species *i* which is capable of growing in monoculture, there is a low enough density of *i* at which the addition of other conspecific individuals necessarily has a beneficial effect on population growth (Figure 1), even if this threshold density is infinitesimal. Setting *α*_*ii*_ to 0 instead means a species will eventually go extinct in isolation. That *β*_*ii*_ must be positive means that above a certain population size of the species, which is determined by the relative strength of *α*_*ii*_ and *β*_*ii*_, addition of conspecific individuals will instead have a negative effect on population growth rate. This allows species *i* to reach a positive equilibrium abundance in isolation. Setting *β*_*ii*_ to 0 means there is no positive stable abundance for *i* in isolation and it will instead continue to grow indefinitely. To summarise, when *N*_*i*_ is low, *α*_*ii*_ drives growth and allows the population to increase. At larger values of *N*_*i*_, the effect of *β*_*ii*_ becomes stronger and limits growth (Fig. 1.B). We note that species that cannot grow in isolation, such as predators, do not need to have a positive *α*_*ii*_. They can instead be modelled with a positive *α*_*ij*_ between predator *i* and prey *j* which will ensure that their population can grow from rare at low densities.

**Figure 1:**
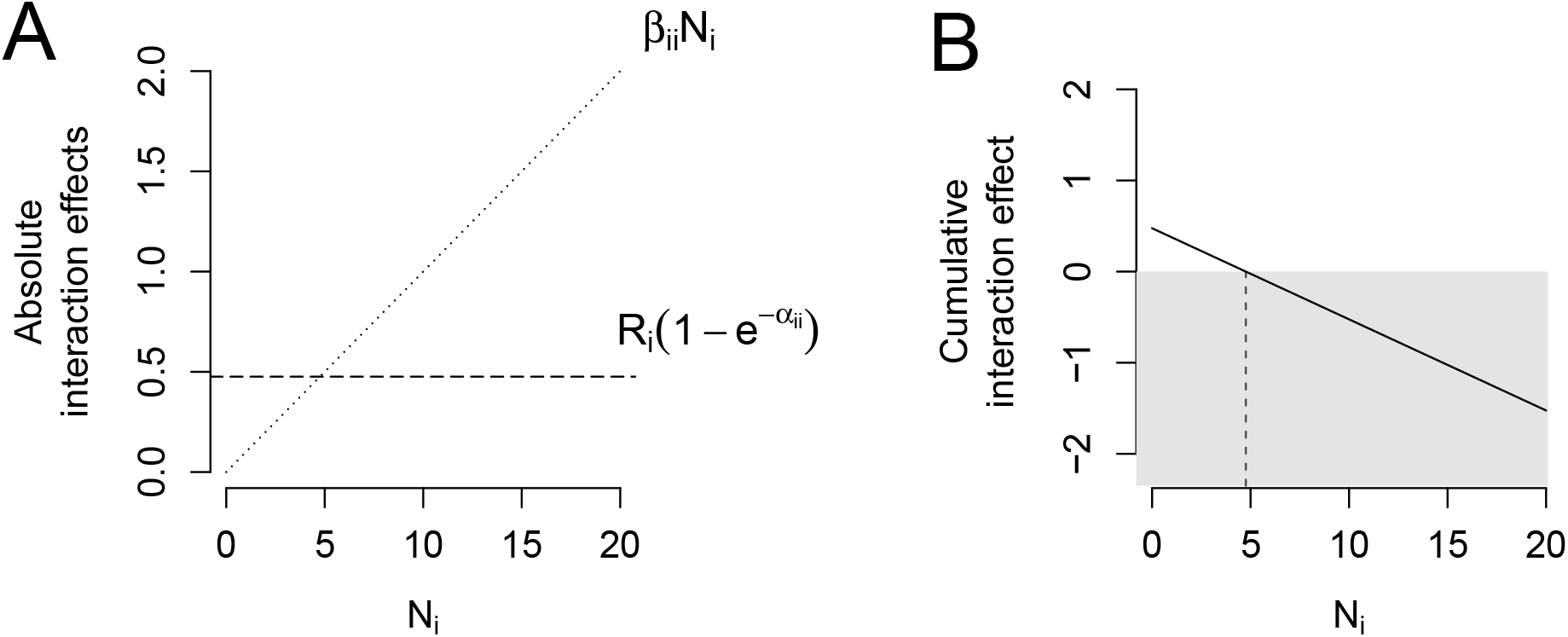
Interactions can vary as a function of density (x-axis), here illustrated for a single population. **A** shows the absolute effects of intraspecific beneficial interactions (dashed line) and intraspecific harmful interactions (dotted line) on *N*_*i*_ as the density of *i* increases. Whilst the effect of beneficial interactions remain constant, the effect of harmful interactions increases with density. **B** shows how taken together, this results in positive density-dependence (white background) at low densities of *i*, the strength of which decreases with density and becomes negative (grey background) at the threshold density where 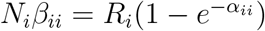 (vertical dashed line).

### 2.2 Relative strength of beneficial and harmful interactions

The *α*_*i\**_ and *β*_*ij*_ parameters describing beneficial and harmful interactions thus have differing effects on model behaviour, and their relative strength varies as a function of population density. This interplay arises from beneficial interactions being placed within an exponent and harmful interactions being placed in a multiplicative formula, justifying our choice of model construction. Similar behaviour as described in the single population case can be extended to multispecies cases where a species *j* has both beneficial and harmful effects on species *i* (Figure 2). Assuming that the density *N*_*i*_ of species *i* is kept constant, the absolute effect of beneficial interactions increases but more and more slowly as the density *N*_*j*_ of species *j* increases whereas the absolute effect of harmful interactions increases linearly with *N*_*j*_ (Fig 2.A). This results in an overall positive effect of *j* on *i* at low densities of *j*, which weakens and eventually becomes negative as *N*_*j*_ increases. Entries in both *α* and *β* can thus be filled to create a threshold at which the overall effect of an interacting species switches from beneficial to harmful. Note that unlike the single-species case where beneficial interactions are constant and harmful interactions are density-dependent, in the multispecies case both interspecific beneficial and harmful interactions are density-dependent but respond at different rates to changes in density.

**Figure 2:**
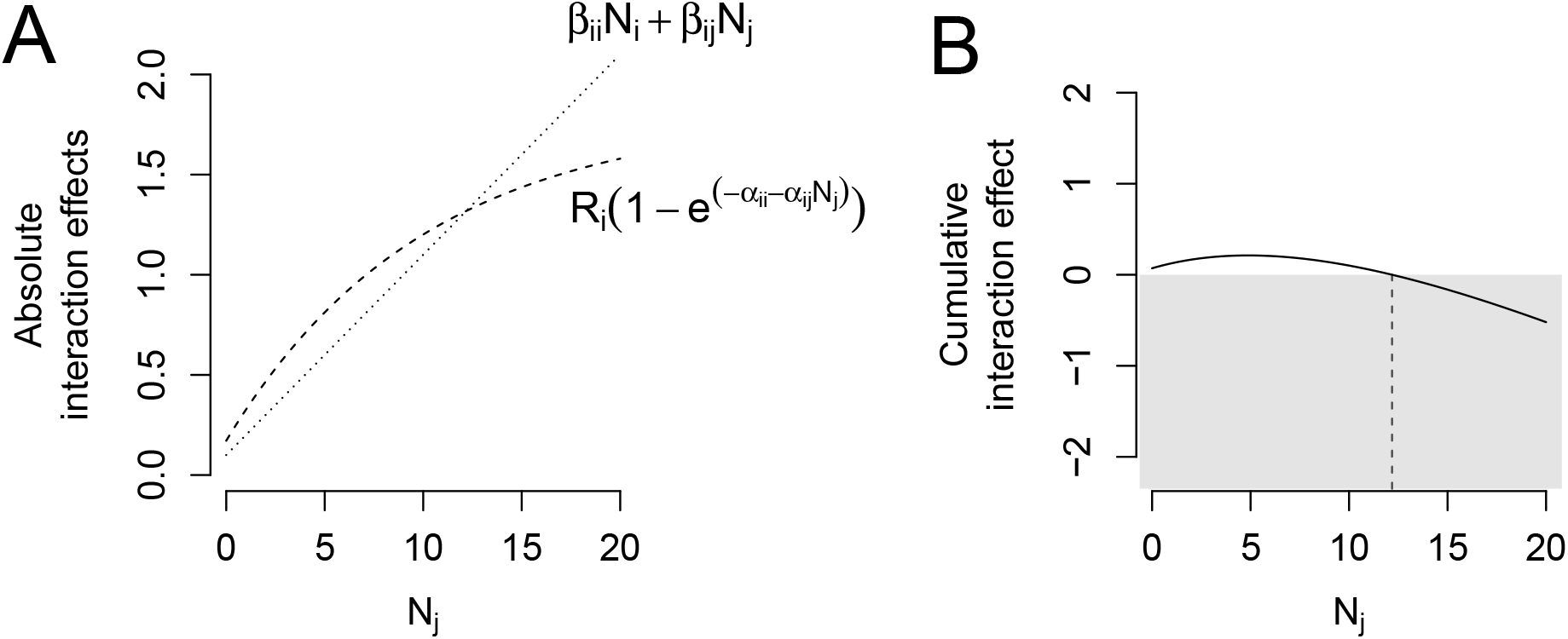
The relative strengths of beneficial and harmful interactions (y-axis) can vary as a function of density, here illustrated for two interacting populations *i* and *j*. Whilst the density *N*_*i*_ of species *i* is kept constant, the density *N*_*j*_ of species *j* is allowed to vary (x-axis). **A** shows how the absolute effects of beneficial (dashed line) and harmful interactions (dotted line) on *N*_*i*_ vary as the density of *j* increases. **B** shows how taken together, the overall effect of species *j* on *N*_*i*_ is positive at low densities of *j* (white background), decreases as *N*_*j*_ increases and becomes negative (grey background) at the threshold density where 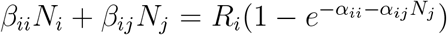 (vertical dashed line). For constant *N*_*i*_, this equation has no analytical solution for *N*_*j*_, so the threshold density of *N*_*i*_ was estimated numerically with the *nleqslv* package (Hasselman 2023).

We emphasise that this switch from an overall beneficial to overall harmful effect remains simply an optional feature of the model and is not strictly required: species can have uniquely beneficial or uniquely harmful effects by simply setting the appropriate *β*_*ij*_ or *α*_*ij*_ parameter to 0, respectively. For example if the presence of species *j* always benefits species *i* (e.g. pollination), *α*_*ij*_ will be positive and *β*_*ij*_ will be equal to 0. As beneficial interactions are included in an exponent term, this automatically creates a saturating effect at high densities, preventing unbounded population growth due to beneficial interactions and removing the need for further saturating parameters when modelling mutualistic systems.

### 2.3 Equilibrium solution for multiple populations

Our model construction means that the equilibrium abundances of multiple interacting species cannot be derived analytically and must instead be solved numerically, for example with the *nleqslv* package in R (Hasselman 2023). We provide the necessary equations to do so in Box 1. These equations also provide insights into the behaviour and stability of our model. In particular, Equation 7 (Box 1) shows that the stability of the equilibrium solution increases when harmful intraspecific and interspecific interactions increase in strength. Conversely, stability decreases as the strength of beneficial interactions increase. Stability can still, however, be ensured despite strong intraspecific and interspecific beneficial interactions as long as the maximum growth rate is sufficiently low. Our model thus presents similar behaviour to GLV models in regards to stability where harmful interactions are concerned, but with the added benefit of also allowing for stable equilibria when beneficial interactions are strong.

#### Box 1

**Equilibrium and stability in the case of multiple interacting species.**

In a multispecies case with several populations interacting with one another, the steady-state solution for Eq. 1 can be found by setting its left-hand side equal to 0. Assuming that the equilibrium abundances of each population *N*_*i*_* *>* 0, we get:

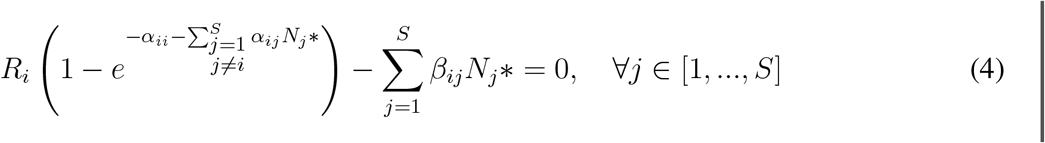

As Eq. 4 are non linear, they cannot be solved analytically. We propose a numerical approach to solve these equations, using Newton’s method. Equation 4 can now be rewritten using matrices. Let *R* be a vector of length *S* containing the maximum growth rates for each species [*R*_1_, …, *R*_*S*_], *A* be the *S* × *S* matrix whose elements are *α*_*ij*_ and *B* the *S* × *S* matrix whose elements are *β*_*i*_*j*. We define *P* as the vector of diagonal elements of *A* such that *P* = [*α*_11_, …, *α*_*SS*_] and *M* a matrix of the same size as *A* where *M*_*ij*_ = 0 if *i* = *j* and *M*_*ij*_ = *α*_*ij*_ if *i* ≠ *j*. In other words, *P* is the vector of intraspecific *α*_*ii*_ whereas *M* contains all interspecific *α*_*ij*_, *i*≠ *j*. Lastly, *N* * is defined as the vector of length *S* of equilibrium abundances with *N* * = [*N*_1_*, …, *N*_*S*_*]. Eq. 4 then becomes:

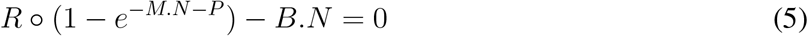

where 1 and 0 are vectors of length *S* and containing all 1 or all 0 elements, respectively. The exponential function is taken elementwise. The symbol ◦ represents the element-wise (Hadamard) product, whereby each element *i* of *R* is multiplied by the *i*’th element of the vectors resulting from the operations within the parenthesis. We can then find the vector of equilibrium abundances which satisfies Equation 5 numerically, for example using Newton’s method with the R function *nleqslv* from the package of the same name (Hasselman 2023). Newton’s method is an iterative numerical technique which can approximate solutions to systems of nonlinear equations by starting with an initial guess and refining it until it converges. This method can find the solution to Eq. 5 provided we supply it with the derivative of the equation to help determine the direction and magnitude of the update in each iteration. This is given by the Jacobian matrix of partial derivatives of Eq. 5 with respect to *N*, 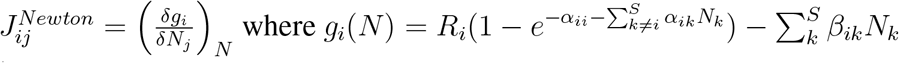. Here *k* is a silent variable to sum all column indexes different from *i*, while *j* denotes the current column index. The Jacobian for Newton’s method is thus defined as:

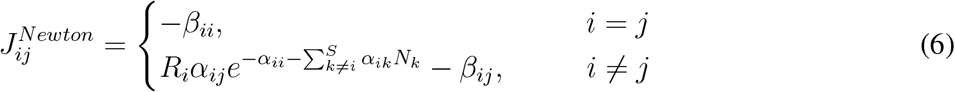

The *nleqslv* function as well as other numerical solvers will return the vector of equilibrium abundances *N* _*i*_*, which is deemed feasible if all equilibrium abundances *N*_*i*_* are positive.

After finding the steady-state solution *N* ^*\**^, we can analyse it for stability to determine whether the system will return to or diverge from equilibrium for small perturbations around the steady state. The Lyapunov stability of the steady-state solution can be verified by calculating the eigenvalues of the Jacobian matrix *J* at equilibrium 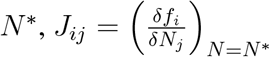 where 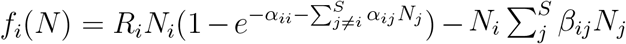. This is a different type of Jacobian matrix to the one defined in Equation 6, because *f*_*i*_(*N*) = *Ng*_*i*_(*N*) and this second Jacobian matrix is only evaluated at *N* * rather than for a variety of *N*. At the steady state *N* * which satisfies Eq. 5, the diagonal and non-diagonal elements of *J* are given by:

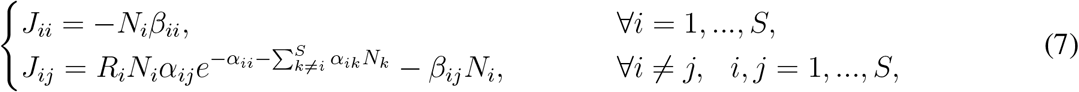

For the system to be deemed stable, the real part of each eigenvalue of *J* must be negative, ensuring that populations will return to their equilibrium values after small perturbations.

## 3 Model behaviour for two-species scenarios

We illustrate the generality of our model and its ability to reproduce both expected and complex ecological patterns by applying it to three simulated scenarios that consist of (1) a mutualistic system, (2) a competitive system, and (3) a predator-prey system. We limit ourselves to two-species systems, firstly to keep the results tractable and secondly, to clearly identify the consequences of varying different model parameters (later, in the case study, we demonstrate the model on a more realistic system). Given the breadth of interaction outcomes our model can potentially describe, simulating every ecological scenario is computationally and time prohibitive. Our two-species scenarios were selected to serve as guidelines for expected model behaviour and benchmark the ‘extremes’ of ecological interaction outcomes (mutually beneficial, mutually harmful, and antagonistic, for scenario one, two and three respectively). Expectations for model behaviour with variable interaction outcomes, such as interactions which switch from beneficial to harmful depending on density, and for more complex systems (i.e. more species and/or different types of interactions) can be built up from the baselines described here and are likely to present more complex dynamics.

Simulations were run in R v.4.3.2 (R Core Team 2023) using the *ode solve()* function from the EEMtoolbox package (Pascal et al. 2024). We focused on varying the relative strengths of the beneficial *α* and harmful *β* interaction parameters (Table 1) to explore model behaviour of interest. Maximum growth rates were set to 1 or 10 depending on trophic level and initial abundances were set to 1. Models were run for 50 years with a resolution of 0.1 years.

**Table 1:** Maximum growth rate and interaction parameter values for each of the three modelled scenarios. Stars (*) indicate parameters with positive, non-zero values that varied across simulations for that scenario. For Scenario 3, species *i* is the the predator and species *j* is the prey.

|  | Maximum<br>growth rate | Beneficial<br>interactions |  | Harmful<br>interactions |  |
| --- | --- | --- | --- | --- | --- |
| Parameters: | $R_i$ | $\alpha_{ii}$ | $\alpha_{ij}$ | $\beta_{ii}$ | $\beta_{ij}$ |
| | $R_j$ | $\alpha_{ji}$ | $\alpha_{jj}$ | $\beta_{ji}$ | $\beta_{jj}$ |
| <u>Scenario 1</u> | 1 | * | * | * | 0 |
| mutualism | 1 | * | 0.1 | 0 | 0.1 |
| <u>Scenario 2</u> | 1 | * | 0 | * | 1 |
| competition | 1 | 0 | 0.1 | * | 0.1 |
| <u>Scenario 3</u> | 1 | 0.01 | 0.8 | 0.1 | 0 |
| predator-prey | 10 | 0 | * | 1 | 0.5 |

### 3.1 Scenario 1: Two-species mutualistic system

In this scenario, species *i* and *j* have purely beneficial effects on one another. Interspecific *α* parameters are thus positive and interspecific *β* parameters are set to 0 (Table 1). Equilibrium abundances *N*_*i*_* and *N*_*j*_* increased with the strength of beneficial interactions and decreased with the strength of harmful interactions (Figure 3). Because the equilibrium abundance of one species is affected by the equilibrium abundance of the other (Box 1, Eq. 4), increasing any of the beneficial interaction strengths (i.e. any *α*) increases both *N*_*i*_* and *N*_*j*_*. This positive interdependence can lead to population explosions in many GLVMs. In our model however, the equilibrium abundances of species *i* and *j* cannot go beyond the maximum set by 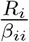 and 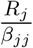, respectively, regardless of how strong beneficial interactions may be.

**Figure 3:**
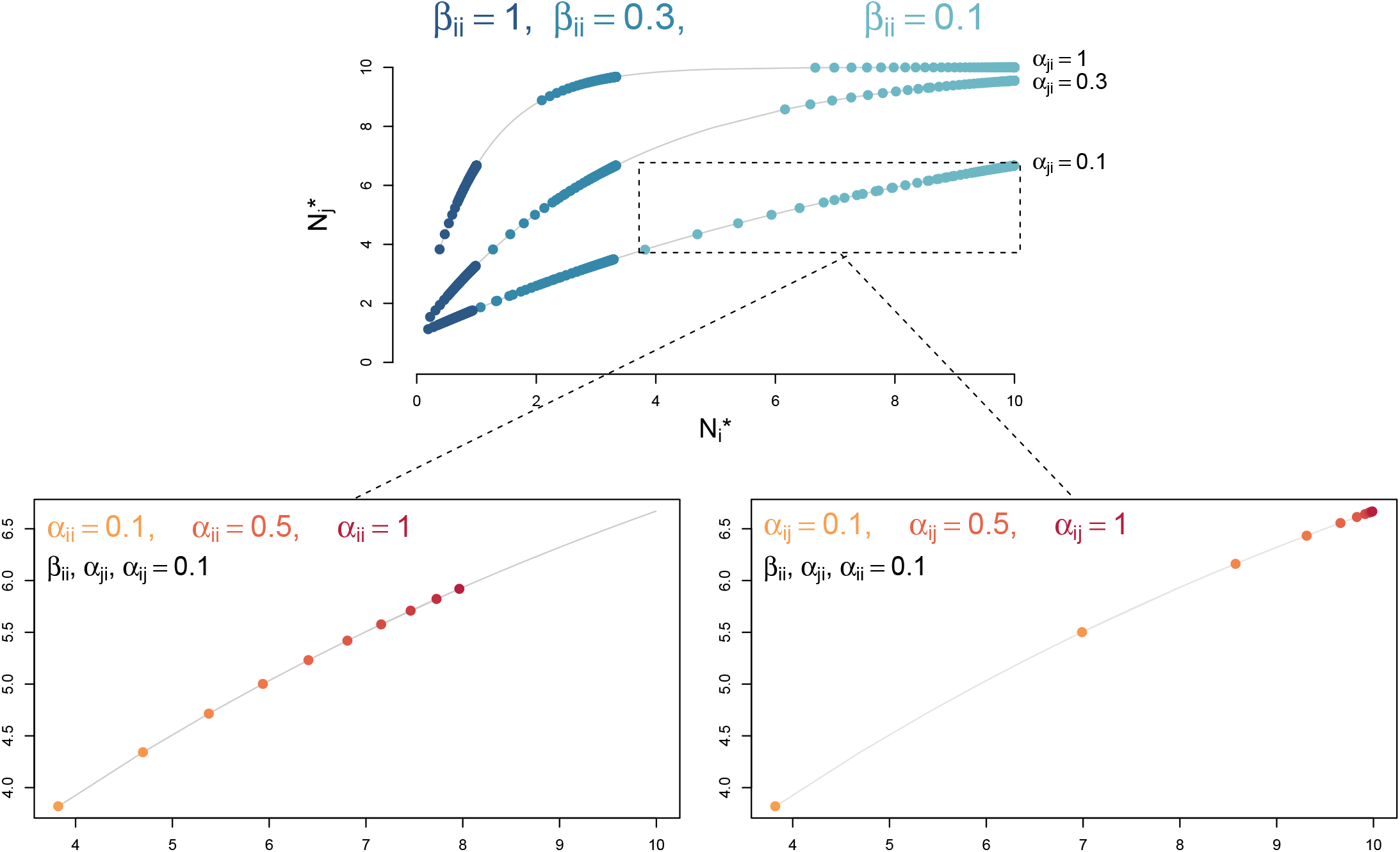
Final equilibrium abundances (x and y-axis) for a two-species mutualistic system after 50 years. Dots correspond to individual simulated predictions, grey lines show *N*_*i*_* as given by Eq. 4 and predicted (simulated) values of *N*_*j*_*. Above, we show how varying *β*_*ii*_ (in blue) and *α*_*ji*_ (grey lines) modify equilibrium values. Bottom panels illustrate the effects of varying *α*_*ii*_ (left) and *α*_*ij*_ (right) by increments of 0.1 whilst other parameters are kept constant, results shown on a subset of simulations only for clarity.

This means that no parameter can influence the final equilibrium abundance of one species solely, as it will necessarily influence the equilibrium abundance of the other. For instance, increasing intraspecific competition *β*_*ii*_ naturally leads to lower equilibrium abundances for species *i*, but also to slightly lower equilibrium abundances for species *j* (blue dots, Fig. 3). Likewise, whilst increasing *α*_*ji*_ primarily increases *N*_*j*_*, it also has a positive effect on *N*_*i*_* (grey lines, Fig. 3). Increasing the strength of *α*_*ii*_ and *α*_*ij*_ also increases the final equilibrium abundances of both species (bottom panels, Fig. 3).

### 3.2 Scenario 2: Two-species competitive system

In the competitive scenario, species *i* and *j* have purely harmful effects on one another. Harmful interspecific *β* parameters are thus positive and beneficial interspecific *α* parameters are set to 0 (Table 1). Increasing the competitive effect of *i* on *j, β*_*ji*_, leads to lower equilibrium abundances for species *j* (grey lines, Figure 4) and sometimes extinction. On the other hand, increasing intraspe-cific competition *β*_*ii*_ leads to much lower equilibrium abundances for *i* and higher equilibrium abundances for *j* (dark blue dots, Fig 4). Even in a predominantly competitive system, our model construction still requires that intraspecific beneficial interactions *α*_*ii*_ and *α*_*jj*_ must be positive and non-zero for both species to be capable of increasing in abundance. The effect of intraspecific beneficial interactions in this competitive scernario is illustrated in the right-hand panel of Figure 4: when all other parameters are kept equal, higher values of *α*_*ii*_ means that species *i* can reach higher equilibrium abundances and that species *j* is more likely to be competitively excluded, compared to when *α*_*ii*_ is low.

**Figure 4:**
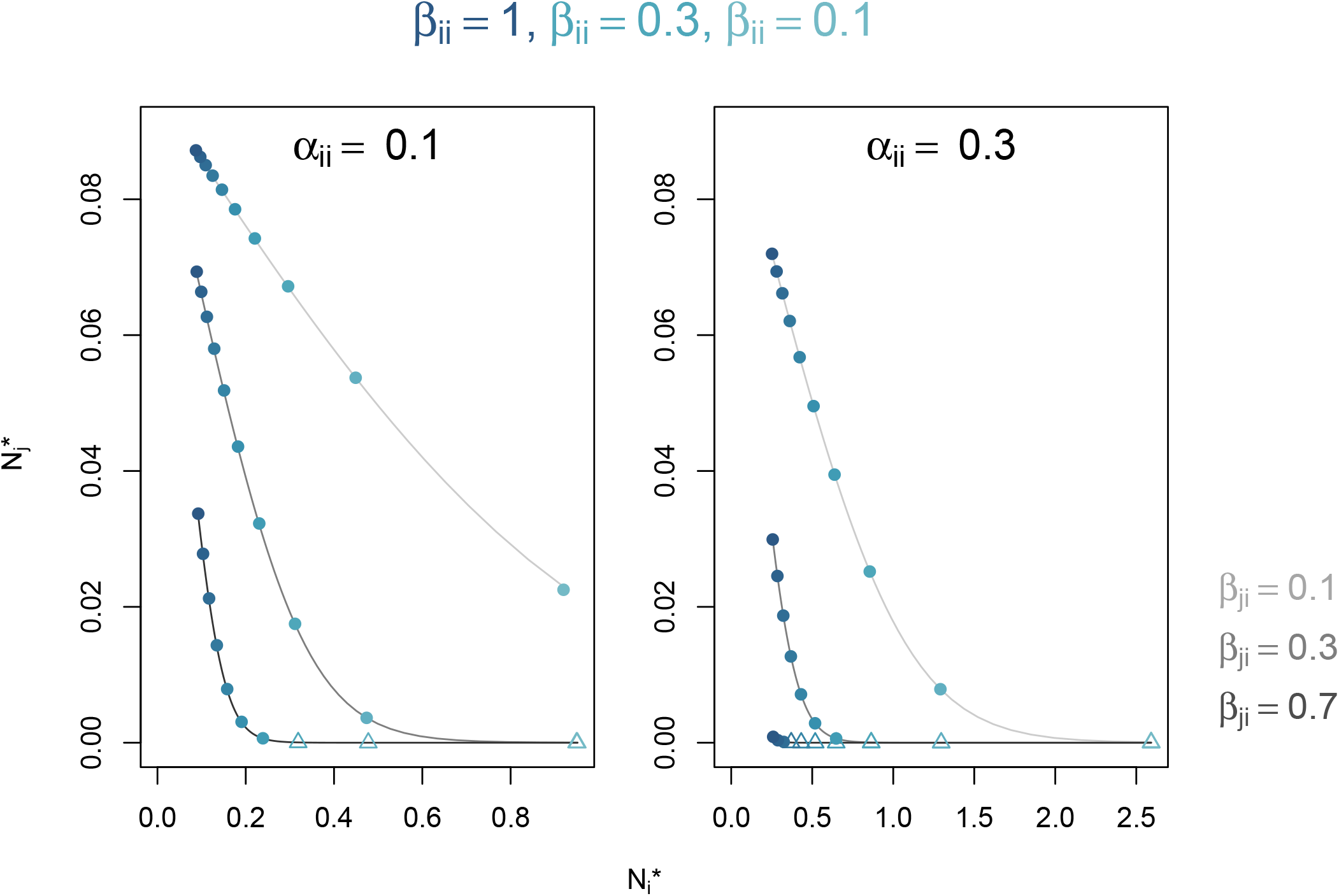
Final equilibrium abundances (x and y-axis) for a two-species competitive system after 50 years. Dots correspond to individual simulated predictions, with darker colours corresponding to higher values of *β*_*ii*_. Triangles indicate simulations where *N*_*j*_* has fallen below 10^*−*5^. Grey lines show *N*_*i*_* as given by Eq. 4 and predicted (simulated) values of *N*_*j*_*. Darker grey lines correspond to higher values of *β*_*ji*_. Left and right panels show simulations under lower and higher values of *α*_*ii*_, respectively. Note the difference in scale of the x-axis for each panel.

### 3.3 Scenario 3: Two-species predator-prey system

In our third scenario, species *i* preys upon species *j*. Here we focus on varying the population growth rate of prey species *j* by manipulating *α*_*jj*_, the strength of prey-prey beneficial interactions. The prey species receives no other beneficial interaction effects and we set the maximum intrinsic growth rate of the prey *R*_*j*_ to be ten times higher than that of the predator *R*_*i*_ (Table 1). A point of distinction between our model and GLVMs is the inclusion of beneficial interactions *α* in an exponent, and as beneficial interactions weaken our model structure begins to resemble that of a GLVM more closely. This is illustrated on the left-hand side of Figure 5: by setting *α*_*jj*_ to a low value, our model can recreate the classic oscillatory dynamics of GLVMs. When *α*_*jj*_ increases (right-hand side, Fig. 5), our model instead exhibits dampened dynamics as despite such strong intraspecific facilitation, prey growth rates remain constrained by the maximum growth rate parameter *R*_*j*_. Whilst oscillatory dynamics can be regained by changing various parameters (and not just *α*_*jj*_), the point remains that in our model, population growth rates are constrained by *R*_*i*_ and *R*_*j*_ which limits the likelihood of population explosions and/or unrealistic growth rates, regardless of how strong species interactions may be.

**Figure 5:**
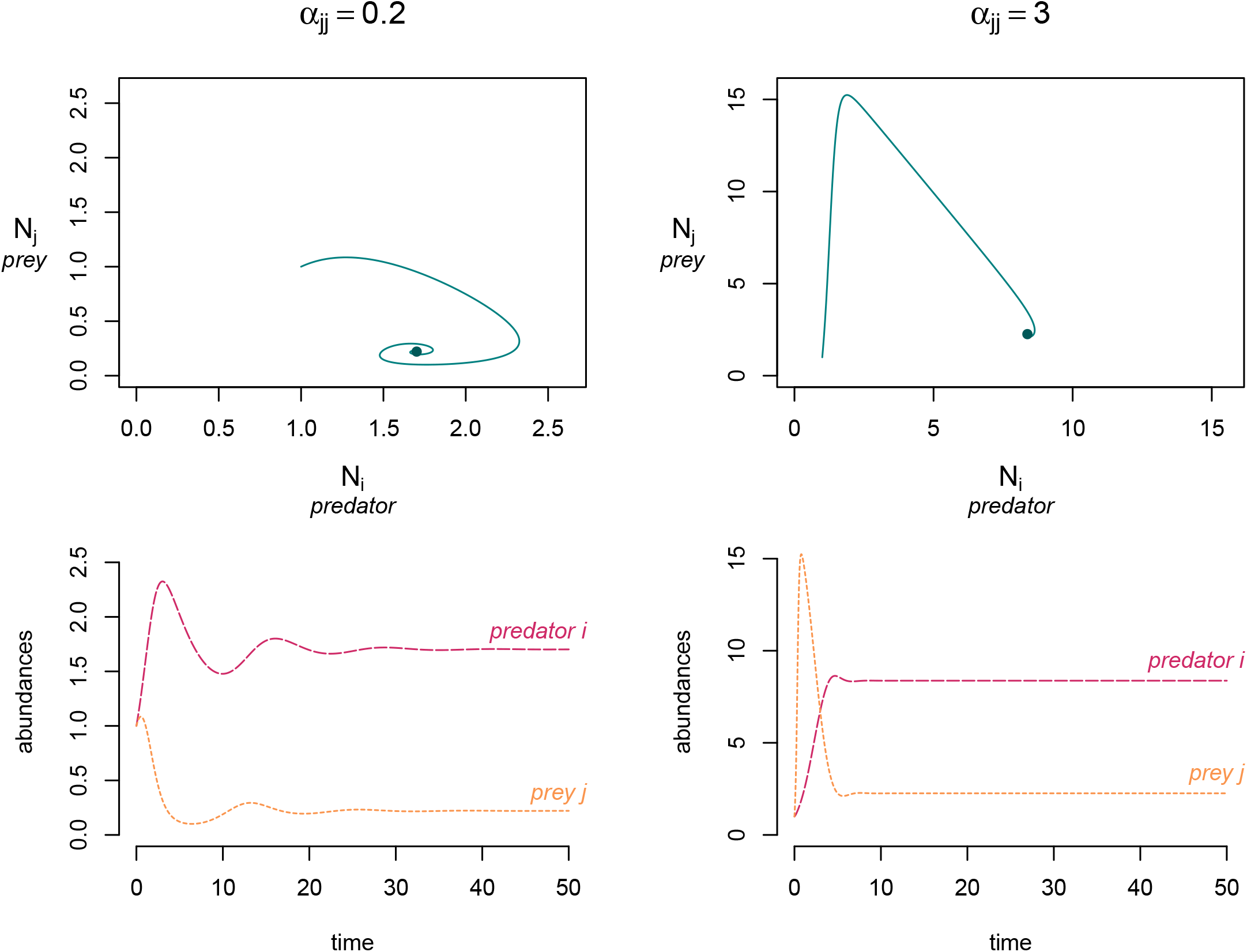
Results from two simulations (left and right) of a predator-prey system after 50 years. Top row shows the phase plane diagrams of predator *i* and prey *j* abundances through time, with the final end points indicated by a blue dot. Bottom row shows the simulated time series, with predator abundances in red and prey abundances in yellow. Note the different scales for abundance between the left and right columns.

## 4 Setting plausible model parameters for empirical applications

A key motivation in the development of our model was to make the parameters more directly relatable to the kind of data that empiricists and other end-users have access to in real-world man-agement scenarios, where information is often sparse. The biggest advantage our model presents in that regard is the use of a maximum growth rate *R*_*i*_, which can be estimated from a wide variety of sources as we illustrate next. We describe the process of setting empirically-informed *R*_*i*_ parameters for a case-study of red-tailed tropicbirds (*Phaethon rubricauda*) from Christmas Island, Australia, which are predated upon by invasive black rats and cats (Plein et al. 2022). Our model also requires the setting of beneficial *α* and harmful *β* interaction parameters. Setting interaction parameters based on empirical data presents a more difficult problem for both our model and others (e.g. Laska and Wootton 1998; Berlow et al. 2004). Here interaction parameters are instead randomly sampled from prior distributions, whose range is constrained based on estimates of carrying capacity.

Christmas Island is a 135*km*^2^ locality situated in the Indian Ocean, with difficult terrain consisting predominantly of tropical rainforest and an inaccessible shoreline making the monitoring of wildlife populations challenging. Whilst the invasive cats and rats both predate on nesting tropicbirds, cats also hunt rats. Thus management strategies which decrease the cat population may also release rats from predation pressure, with unintended consequences for the tropicbird population and other native species. Plein et al. (2022) applied a bio-energetic predator-prey model to investigate how different combinations of rat and cat abundances might affect population trajectories of the red-tailed tropicbirds. We use our model here to ask whether reducing the cat population will have harmful or beneficial effects on the red-tailed tropicbird population. We begin by searching the literature to assign values to the maximum intrinsic growth rate *R*_*i*_, using the subscripts *i* = *t, i* = *c* and *i* = *r* for tropicbirds, cats, and rats, respectively. We then use carrying capacity estimates to inform the relative strength of intraspecific beneficial interactions *α*_*ii*_ and intraspecific harmful interactions *β*_*ii*_ before sampling the remaining unknown interspecific interaction parameters.

### 4.1 Red-tailed tropicbird parameters

As detailed by Plein et al. (2022), red-tailed tropicbirds reach reproductive maturity at three years old (del Hoyo et al., 1992) and mate monogamously for life, with each breeding pair producing one egg per breeding season for up to 16 years (Schreiber & Schreiber 1993, *situ* Plein et al.2022). Given that we are interested in the *maximum* growth rate *R*_*t*_ that is achieved under optimal conditions, we assume the most generous estimates and calculate the number of eggs per bird over its reproductive lifespan divided by total lifespan: *R*_*t*_ = (0.5 × 16) ÷ 19 = 0.421 yr^*−*1^.

Carrying capacity in the absence of cats and rats is difficult to estimate given all three species have co-occurred on Christams Island for over 100 years, but historical estimates range from 1440 to 2000 breeding pairs (James et al 2014, Stokes 1988, *situ* Plein et al. 2022). We assume these numbers are significant underestimates and set the single-species carrying capacity 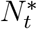 to 10000 birds. We can now relate both of these numbers to Eq. 3 such that 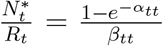, substituting our estimates into the left-hand side of the equation. This give us the relationship between intraspecific beneficial interactions *α*_*tt*_ and intraspecific harmful interactions *β*_*tt*_, though their exact values remain unknown. We apply the following thought-process to constrain the limits of the parameter space we have to sample to determine the intraspecific interaction parameters. The term 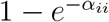 can be thought of as the percentage of the maximum growth rate *R*_*i*_ which is realised in the absence of other species in the model, prior to any population decline (which is encapsulated by the *β* parameters, see Eq. 1). Given that red-tailed tropicbirds rely on neither cats nor rats to grow their population, we assume that in the absence of these invasive species the tropicbird population achieves 80% to 100% of it’s maximum growth rate in any given year. We thus sample 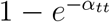 from a uniform distribution of range (0.8, 1) and input these values into Eq. 3 to arrive at estimates of *β*_*tt*_.

### 4.2 Cat and rat parameters

Demographic parameters for cats and rats on Christmas Island are unavailable in Plein et al. (2022). To ensure we still set biologically reasonable estimates, we select studies of feral cat populations in mainland USA (Nutter et al. 2004; Foley et al. 2005) and on the sub-Antartic Marion Island (Van Aarde 1983), as well as a review of invasive rat population biology on tropical islands (Harper and Bunbury 2015). The following numbers are not intended to be foolproof and a more extensive review of the literature would likely refine their accuracy; we opt for brevity to succinctly illustrate the thought-process.

Cats can have their first litter as early as six months old, with up to six kittens per litter and two litters per year (Nutter et al. 2004; Van Aarde 1983). Whilst three litters is possible, this is linked to 100% mortality of the first or second litter (Nutter et al. 2004). The lifespan of feral cats is much shorter than domestic cats, and Van Aarde (1983) reports a 100% mortality rate for feral cats over 8 years old. We thus assume a female cat can produce one litter of 6 kittens in her first year and two litters of 6 kittens each year in the following 7 years, setting the maximum growth rate *R*_*c*_ to 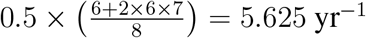.

Unfortunately the Van Aarde (1983) study finds the Marion Island cat population is still increasing and thus inappropriate for setting equilibrium density, but Foley et al. (2005) provide carrying capacity estimates of 210325 and 19323 feral cats in San Diego County (11720km^2^) and Alachua County (2510km^2^) respectively, which gives us density estimates of 17.946 and 7.7 cats per km^2^ respectively. Taking the average, we roughly extrapolate a density of 12.822 cats per km^2^ over the 135km^2^ of Christmas Island, resulting in a single-species carrying capacity 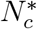 of 1731 cats. We set *α*_*cc*_ and *β*_*cc*_ following the same process as for tropicbirds, but this time we assume that in the absence of both rats and birds, the cat population does not have access to its preferred food sources and only achieves 20% to 40% of its maximum growth rate. We thus sample 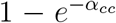 from a uniform distribution of range (0.2, 0.4) from which we then determine *β*_*cc*_.

Black rats *Rattus rattus* typically have higher population densities and number of offspring on tropical islands than temperate islands. Harper and Bunbury (2015) report an average litter size of 4.47 on tropical islands, with a maximum of 6.3 (Davis 1953). We assume rats live for one year. Juveniles can reproduce within three to five months of their birth and females have been reported to produce up to 5 litters per year, we assume a maximum of 4 litters per year to account for the pre-reproductive stage. We thus calculate the maximum growth rate of rats to be *R*_*r*_ = (0.5 × 4 × 6.3) = 12.6 yr^*−*1^.

Black rat population densities are incredibly variable between tropical islands but follow rainfall patterns, so we use an estimated maximum density of 38ha^*−*1^ from Isabel Island which has a similar climate to Christmas Island (Harper and Bunbury 2015). This gives a single-species carrying capacity 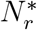 of 513000 rats. Rat diets appear to depend far less on birds than do cat diets (Plein et al. 2022) so we assume that in the absence of both cats and birds, the black rat population on Christmas Island achieves 50% to 70% of its maximum growth rate and sample 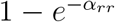 from a uniform distribution of range (0.5, 0.7) to arrive at *β*_*rr*_.

### 4.3 Simulations and results

In our case study, the strength of beneficial and harmful interactions between cats, rats and tropicbirds is unknown. We use simulations to vary the strength of interspecific interactions between all three species, following the interaction structure described in Figure 6. We randomly sample *α*_*ij*_ values from a uniform distribution with a range of 0.1−2 and *β*_*ij*_ values from a uniform distribution with a range of 1 × 10^*−*5^ − 1 × 10^*−*3^ to match the range of values assigned to *α*_*ii*_ and *β*_*ii*_, respectively. We then apply the *nleqslv* function (Hasselman 2023) as described in Box 1 to determine equilibrium abundances for our sets of empirically-derived and simulated parameters, discarding those sets which resulted in equilibrium abundances of 1 or below for any of the three species. Of those sets where all species achieve equilibrium abundances above 1, we randomly selected 1000 sets for which to simulate cat removal. We use the *ode solve* function from the EEMToolbox package (Pascal et al. 2024) to simulate cat eradication from the system, setting cat initial abundance to 0 and initial abundances of birds and rats to their respective equilibrium values for each of the 1000 sets of parameters, running the model for 20 years. We find that red-tailed tropicbird abundance goes extinct across all 1000 simulations, suggesting that cat predation strongly mitigates the potential damage caused by rats on the native seabirds.

**Figure 6:**
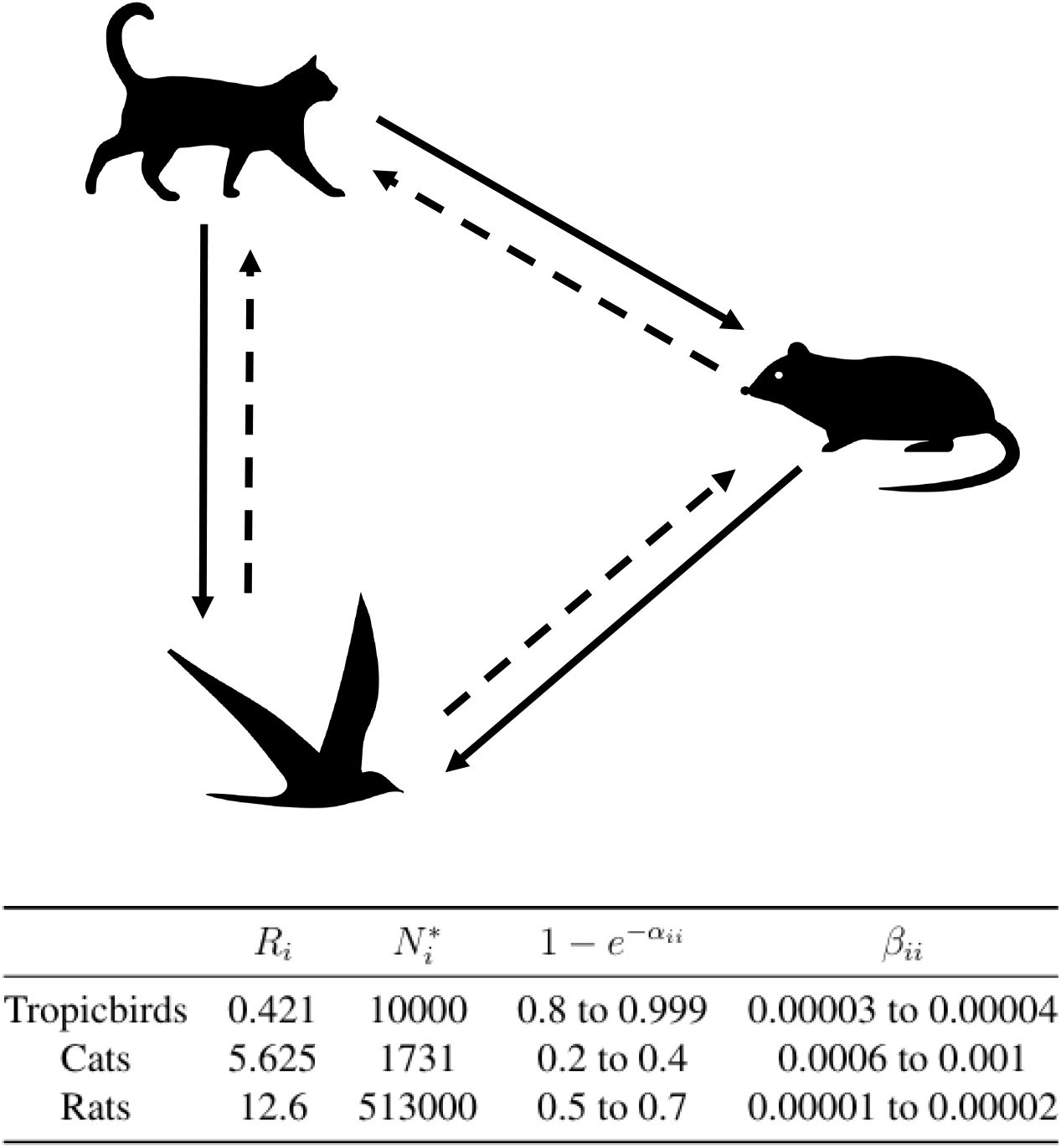
Above, the interaction matrix structure between cats, rats and red-tailed tropicbirds on Christmas Island. Interspecific harmful *β* parameters are represented as solid arrows and interspecific beneficial *α* parameters are dashed arrows. Arrows point towards the species receiving the effect of the interaction. Below, a table summarising the growth rate and single-species carrying capacities assigned to each species, as well as the range of values from which the intraspecific interaction parameters are drawn.

## 5 Discussion

Our paper presents a new generaliseable population dynamic model developed for application to data-poor systems with variable interaction outcomes. This model describes the dynamics of both beneficial and harmful interaction outcomes, allowing the inclusion of a wide range of trophic and non-trophic interactions together such as competition, mutualism, and predation, all within the same model equation. Additionally, parameterisation of the growth rates requires relatively little data, as illustrated by our case study. These advantages aim to maximise our model’s usefulness to a broad range of ecological and management contexts, where natural communities are often structured by variable and diverse sets of interactions and data availability fluctuates widely. We have focused our presentation of this paper towards developing intuition for our newly introduced parameters and their behaviour in an effort to further increase ease-of-use and accessibility of the model.

Our model introduces a maximum intrinsic growth rate *R* that naturally constrains population growth below its value. This has two advantages. First, the inclusion of beneficial interactions such as mutualism becomes more straightforward as there is no risk of unbounded population increases or overexplosions: once *R* is set within reasonable bounds, population growth remains constrained regardless of the interaction structure. Secondly, our model growth rates are easier to parameterise than the intrinsic growth rates of GLVMs, as it is arguably simpler to estimate a maximum value from empirical data than a mean. To clarify, the *R* parameter can be interpreted as the maximum rate a population grows under optimal conditions. *R* should thus almost always be higher than a growth rate estimated from field data, where competitors, predators, and limited resources make conditions less than ideal. Instead, *R* can be set based on records of reproductive output such as maximum clutch or litter size, estimates which can be drawn from a wide variety of sources beyond population dynamics studies such as natural history observations, breeding programs or historical demographic data. This presents in advantage compared to the growth rate parameter *r* used in GLVMs, defined as the low density intrinsic growth rate, which assumes that low abundance coincides with optimal conditions for growth (because intraspecific competition is minimal). Not only is this assumption not always verified (e.g. in the case of facilitation or mutualisms), but natural populations cannot always be observed at low density, often leading to incorrect estimates (Cortés 2016).

Separating out the effects of interactions — traditionally described by a single interaction matrix — into two interaction matrices allows us a great degree of flexibility in how we represent the outcomes of ecological processes mediated by interactions. For example, plant-pollinator mutualisms and intraguild competition between plants both cooccur within the same communities yet are often described with separate equations (e.g. Thébault and Fontaine 2010; Yacine and Loeuille 2022). Both interaction types can now be included under the same equation, within the beneficial and harmful interaction matrices, respectively. The model does not prescribe certain interaction types to particular trophic levels either: we could also include plant-plant facilitation, and competition between pollinators. Because both beneficial and harmful interactions are density-dependent but at different rates, interactions can be modelled to have a beneficial outcome at low density, which gradually becomes harmful as species density increases. While we do not explore this behaviour presently, the dynamics would likely be more complex than the scenarios we described. The two interaction matrices can thus serve as building blocks for more complex, density-dependent interactions.

The presence of two interaction matrices does however, makes it difficult to infer interaction strength values from abundance data (for example through regression [give examples?]). Model applications are thus restricted to systems where the interaction structure is already established (knowledge of which species interact and whether the outcome is beneficial, harmful, or both), and interaction strengths can instead be simulated. As in the case study, different combinations of interaction strengths can be kept or discarded depending whether the outcome matches empirical observations, and model behaviour explored for those sets of parameters only (Baker et al. 2016). A recent R package, *EEMToolbox*, provides this functionality for several population models including the one presented here (Pascal et al. 2024).

We are not the first to suggest a dual interaction matrix to model beneficial and harmful interactions together, though previous studies tend to focus on the theoretical implications of this model construction. Many models of mutualisms include terms for both beneficial and harmful interac-tions and require the addition of second or third-order terms to ‘brake’ population growth when it reaches a certain maximum. Recently, Aguadé-Gorgorió et al. (2024) explore the stability conditions of a community model with an Allee effect and dual interaction matrices where interaction outcomes can also switch from beneficial to harmful. Given the similarity between their model and the one described here, it is likely that their conclusions extend to our case. Nevertheless, a thorough examination of the conditions for feasibility and stability in our model would likely yield novel insights as multiple interaction types are rarely studied simultaneously (Fontaine et al. 2011).

## Acknowledgements

MDB would like to thank Margie Mayfield, Hanlun Liu, Daniel Stouffer and members of the May-field Lab for thoughtful discussions on this project. MDB acknowledges funding from the Botany Foundation. MPA’s contribution was funded by an ARC Discovery Early Career Researcher Award (DE200100683). LVP and MPA acknowledge funding from the ARC SRIEAS Grant SR200100005 Securing Antarctica’s Environmental Future. LVP acknowledges funding from an Australian Research Council Discovery Project (DP200102101), from an Australian Research Council Discovery Early Career Researcher Award (DE200101791) and from the QUT Centre for Data Science for funding this research through the Second Byte Funding Program 2024.

## Supplementary Information

### S1 Relationship to GLV models

In the interest of comparing our novel model with the generalised LV models that are commonly used to model population dynamics, we show here how they are mathematically equivalent at low growth rates, but model behaviour diverges as growth rates increase. We can approximate Eq. 1 using a Taylor series expansion around 0. The expansion for the exponential function is:

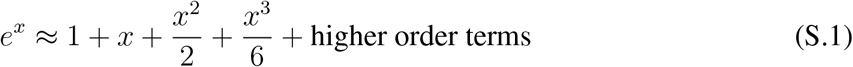

Substituting the first two terms of Eq. S.1 (1 + *x*) and dropping any higher order terms into Eq. 1, we get:

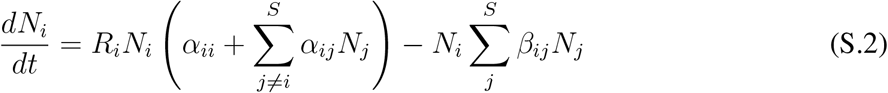

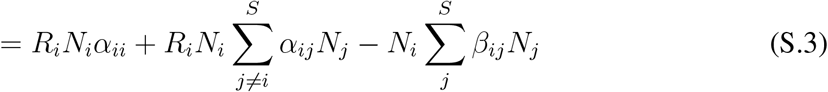

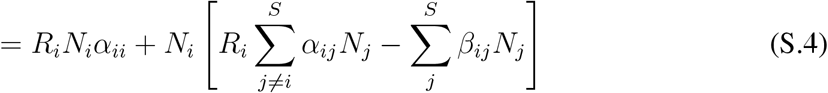

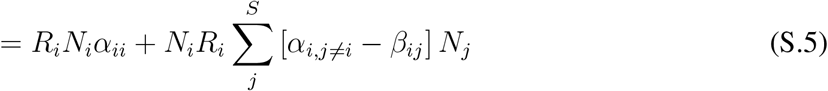

We then define

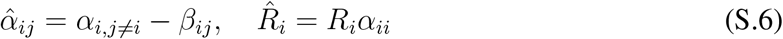

Substituting Eq. S.6 into Eq. S.5, we get

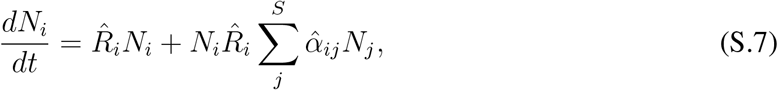

which is the standard GLV form.

This working is only valid while the Taylor series substitution (from Eq. S.1) is a reasonable approximation. That is, while

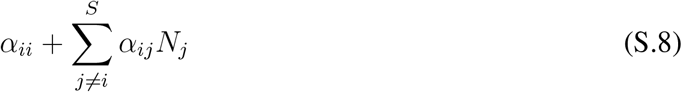

is small, which is the growth term. Hence, at low growth rates, Eq. 1 is equivalent to a generalised LV model. As growth rates increase however, abundance in our model asymptotes such that there is a limit to how fast a population can grow. This behaviour contrasts with GLV’s which allow unbounded population growth, and thus typically require the diagonal elements of the interaction matrix be larger than the off-diagonal elements to prevent runaway population growth.

